# Neurotransmitter circuits of the human language network and their disruption in post-stroke aphasia

**DOI:** 10.64898/2026.05.14.725241

**Authors:** Pedro Nascimento Alves, Isabel Pavão Martins, Chris Rorden, Marcelo L Berthier, Guadalupe Dávila, Alexander Leff, Michel Thiebaut de Schotten, Stephanie J Forkel

## Abstract

Language depends on distributed brain networks whose neurochemical organisation remains largely unknown. Here, we combine large-scale functional MRI, positron-emission tomography–derived receptor and transporter maps, and structural connectomics to characterise the neurotransmitter architecture of the human language network and its disruption in post-stroke aphasia.

We show that language circuits are organised by connection class: cortico-cortical pathways are dominated by serotonergic and glutamatergic signalling, thalamo-cortical projections are predominantly cholinergic, and anterior temporal connections show GABAergic circuit enrichment. In 239 stroke patients, aphasia was associated with distinct neurochemical disruption profiles, with lobar lesions showing postsynaptic serotonergic injury and subcortical aphasia showing preferential presynaptic cholinergic or serotonergic disruption. These findings map the neurochemical architecture of the language network and establish a neurotransmitter circuit-level framework for post-stroke aphasia.

The post-hoc analysis of four pharmacological trials revealed an association between treatment-to-neurotransmitter matching and aphasia improvement, suggesting that personalised pharmacological strategies may enhance behavioural intervention responsiveness.

## 1. Introduction

Language is one of the most distinctive hallmarks of human cognition and behaviour.^1^ The capacity to speak, comprehend, read, and write enables communication and underlies activities ranging from everyday interactions to the most advanced scientific and societal achievements. Disorders of language resulting from acquired brain injury—most commonly stroke—are associated with substantial disability, diminished quality of life, and emotional distress.^2–9^

Historically, post-mortem studies were instrumental in identifying key left-hemispheric regions implicated in aphasia. However, the advent of neuroimaging has revolutionised our understanding of the neural basis of language in vivo. Techniques such as voxel-based lesion-symptom mapping (VLSM), functional MRI (*f*MRI), and diffusion-based tractography have provided new and complementary insights, refining classical models of language processing.^10–21^ Yet, despite these advances, available treatments for aphasia remain insufficient to fully restore language function in a large proportion of cases.^3,8,22^ Notably, several therapeutic approaches, ranging from pharmacological interventions to neuromodulation and speech and language therapy, are thought to exert their effects by modulating neurotransmitter systems. However, the neurochemical organisation of the language network itself remains poorly characterised.

Neurotransmitters represent a fundamental pillar of brain function. Through neurochemical signalling, neurotransmitters orchestrate the dynamics of neuronal networks that support cognition and behaviour.^23–29^ Glutamate and GABA are the main fast-acting excitatory and inhibitory neurotransmitters, respectively, while acetylcholine, dopamine, noradrenaline, and serotonin modulate large-scale brain circuits involved in cognitive and behavioural regulation.^30–33^ The neurons that produce these modulatory neurotransmitters originate in specific nuclei of the brainstem and basal forebrain and project broadly to cortical and subcortical regions, including the thalamus and basal ganglia.^31^

In the context of language, some neurochemical studies have begun to uncover the receptor-level specialisation of language-related cortical regions. A Jülich–Leipzig collaboration combining quantitative autoradiographic receptor mapping with linguistically motivated definitions of language-related cortical regions in post-mortem human brains demonstrated that these regions exhibit distinct neurotransmitter receptor fingerprints compared with non-language areas, particularly in serotonergic 5HT1a and muscarinic acetylcholine receptor densities.^34^ Further evidence implicates GABAergic signalling in language processing, with studies showing a relationship between GABA concentrations in the anterior temporal lobe and semantic processing.^35,36^ Nevertheless, a comprehensive model of the neurotransmitter systems underpinning language remains to be established. Two major limitations may be hindering this progress.

First, in vivo assessment of neurotransmitter circuit integrity remains constrained. Positron Emission Tomography (PET) is the gold standard for mapping neurotransmitter receptors and transporters, but it is impractical and potentially unsafe to repeatedly scan an individual to characterise different receptor systems. Recently, Hansen and colleagues published a normative atlas of receptor and transporter densities based on PET data.^37^ Building on this resource, we combined these maps with tractography data to trace white matter projections associated with specific neurotransmitters, creating a normative atlas of neurotransmitter circuits.^38^ Subsequently, Koch and colleagues independently achieved the same goal using a different methodological approach and obtained comparable results.^39^ We also developed NeuroT-Map, a tool that estimates the degree to which a focal brain lesion disrupts a given neurotransmitter circuit and whether the disruption is predominantly pre- or postsynaptic.^38^

Second, group-average alignment of *f*MRI data based solely on anatomical landmarks often fails to precisely localise small subcortical nuclei involved in language processing. This is due to the low signal contrast, small volume, and substantial inter-individual cytoarchitectonic variability of subcortical structures.^40,41^ This is particularly relevant for language, as subcortical nuclei act as key relays within neurotransmitter systems, and converging evidence from lesion studies and task-based *f*MRI meta-analyses indicates that the thalamus and basal ganglia are essential components of the language network.^42–44^ Functional alignment methods have been shown to enhance anatomical-functional correspondence, particularly in subcortical regions, and may thus improve detection of their involvement in language-related circuits.^23,45–48^

An integrative model of the neurotransmitter systems underlying language would advance our understanding of its neurobiology and may also have direct pharmacological implications. Several clinical trials have investigated the use of neurotransmitter-modulating drugs as adjunctive treatments in post-stroke aphasia. Acetylcholinesterase inhibitors have shown potential to improve naming, phonetic production, and speech fluency, although results on comprehension have been inconsistent.^49–52^ Trials with levodopa and dopamine receptor agonists have yielded similarly inconsistent findings—some showing benefit in verbal fluency and repetition, others reporting no effect.^53–56^ Evidence for serotonin–noradrenaline reuptake inhibitors is more limited, but a recent trial suggested potential efficacy of venlafaxine in patients with subcortical aphasia.^57^ Serotonin-selective reuptake inhibitors were associated with language improvement in two longitudinal studies, but three large trials of fluoxetine found no benefit for patient-reported communication outcomes.^58–61^ Together, these mixed results suggest that variability in treatment response may reflect heterogeneity in the underlying neurochemical disruption across patients, highlighting the need for mechanistic stratification beyond anatomical lesion location alone. The ability to predict which neurotransmitter systems are disrupted in a patient, and whether the damage is predominantly pre- or postsynaptic could be key to guiding personalised pharmacological interventions. Specifically, quantifying the balance between pre- and postsynaptic dysfunction may inform tailored treatment strategies.^38^

In this study, we pursued three objectives. First, we sought to identify the neurotransmitter circuits underlying the language network. Using integrative functional, structural, and neurochemical mapping, we demonstrate that cortico-cortical language pathways are primarily serotonergic and glutamatergic, while acetylcholine-related projections dominate thalamo-cortical tracts, and GABAergic circuits are prominent in anterior temporal regions. Second, we examined whether and how patterns of neurotransmitter circuit disruption differ in post-stroke aphasia. We observed that most cases exhibited predominant serotonergic postsynaptic disruption, whereas subcortical aphasia was associated with a distinct pattern of presynaptic damage to acetylcholine or serotonergic systems. Finally, we evaluated in a post hoc analysis whether neurotransmitter disruption profiles predicted patient response across four clinical trials. Patients whose medication targeted the dominant disrupted neurotransmitter were more likely to show improvement in aphasia outcomes.

## 2. Results

### 2.1. Neurotransmitter circuits of the language network

#### 2.1.1. Cortical and subcortical regions of the language network

To define the cortical regions of the language network, we used the *f*MRI dataset provided by Lipkin and colleagues.^62^ The dataset comprises individual *f*MRI statistical maps from 806 healthy participants performing language tasks (sentence reading versus nonword reading, n = 688; other reading contrasts, n = 105; and listening contrasts, n = 13; see Methods for details). Individual t-maps were thresholded at p<0.05 and activation clusters present in at least 50% of individuals were identified.^62^

The analysis revealed six reproducible cortical clusters: the left lateral temporal cortex, left inferior frontal gyrus, left precentral and middle frontal gyri, left superior frontal gyrus, right lateral temporal cortex, and right cerebellar cortex (Figure 1a).

**Figure 1.**
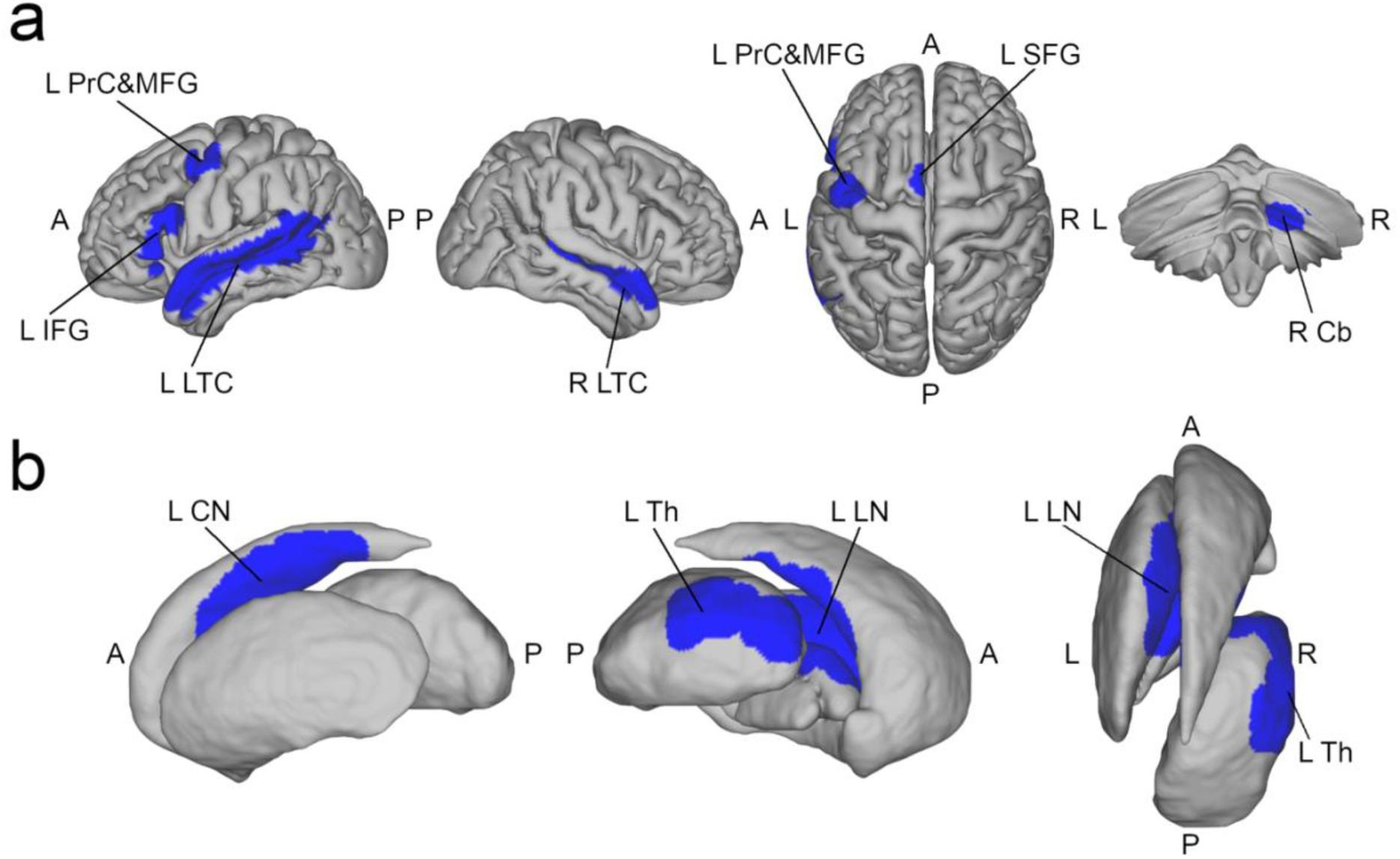
Language clusters identified from the language *f*MRI task. Group-consistent cortical (a) and subcortical (b) regions identified from individual language-task *f*MRI maps (provided by Lipkin et al.^62^). Cortical clusters were defined as voxels present in at least 50% of participants, and subcortical clusters as voxels present in at least 25% of participants, following functional inter-individual alignment. Blue overlays indicate regions consistently engaged during language tasks across individuals. A, anterior; L, left; L CN, left caudate nucleus; L IFG, left inferior frontal gyrus; L LN, left lentiform nucleus; L PrC&MFG, left precentral and middle frontal gyri; L SFG, left superior frontal gyrus; L Th, left thalamus; P, posterior; R, right; R Cb, right cerebellum.

As previously noted, subcortical regions are particularly susceptible to functional misalignment in group analyses based solely on anatomical registration, owing to low signal contrast, small volume, and inter-individual cytoarchitectonic variability.^40,41^ To address this limitation, we applied functional intersubject alignment after anatomical registration and used a 25% voxel-overlap threshold for subcortical cluster identification.^23,45–48^

Two consistent subcortical clusters were identified: one located in the anteromedial region of the left thalamus, and the other in the superior portion of the left lenticular nucleus and the inferior portion of the left caudate nucleus (Figure 1b).

#### 2.1.2. Neurotransmitter white matter projections

To map the neurotransmitter receptor and transporter densities onto the language structural network, we used the Functionnectome.^38,63^ This method projects grey matter voxel values onto white matter based on voxel-wise, weighted probabilities of structural connectivity.^63^ Structural connectivity priors were defined using normative structural probability maps derived from whole-brain deterministic tractography of 7T diffusion-weighted MRIs from 176 healthy participants of the Human Connectome Project. We selected only the tracts connecting at least two language clusters.^64^ PET-derived density maps provided by Hansen and colleagues were used as inputs to the Functionnectome.^37^

The association fibres between language cortical areas were predominantly serotonergic relative to other modulatory neurotransmitter systems (Figure 2a). Specifically, the tracts connecting the left lateral temporal cortex and the left inferior frontal gyrus were particularly rich in postsynaptic 5HT6R and 5HT2aR serotonergic fibres, followed by noradrenergic presynaptic fibres (Figure 2b). The neurochemical profile of the ventral pathway was similar to that of the dorsal pathway (Supplementary Figure 1). The commissural fibres linking the left and right lateral temporal cortices were also mainly serotonergic (Figure 2a).

**Figure 2.**
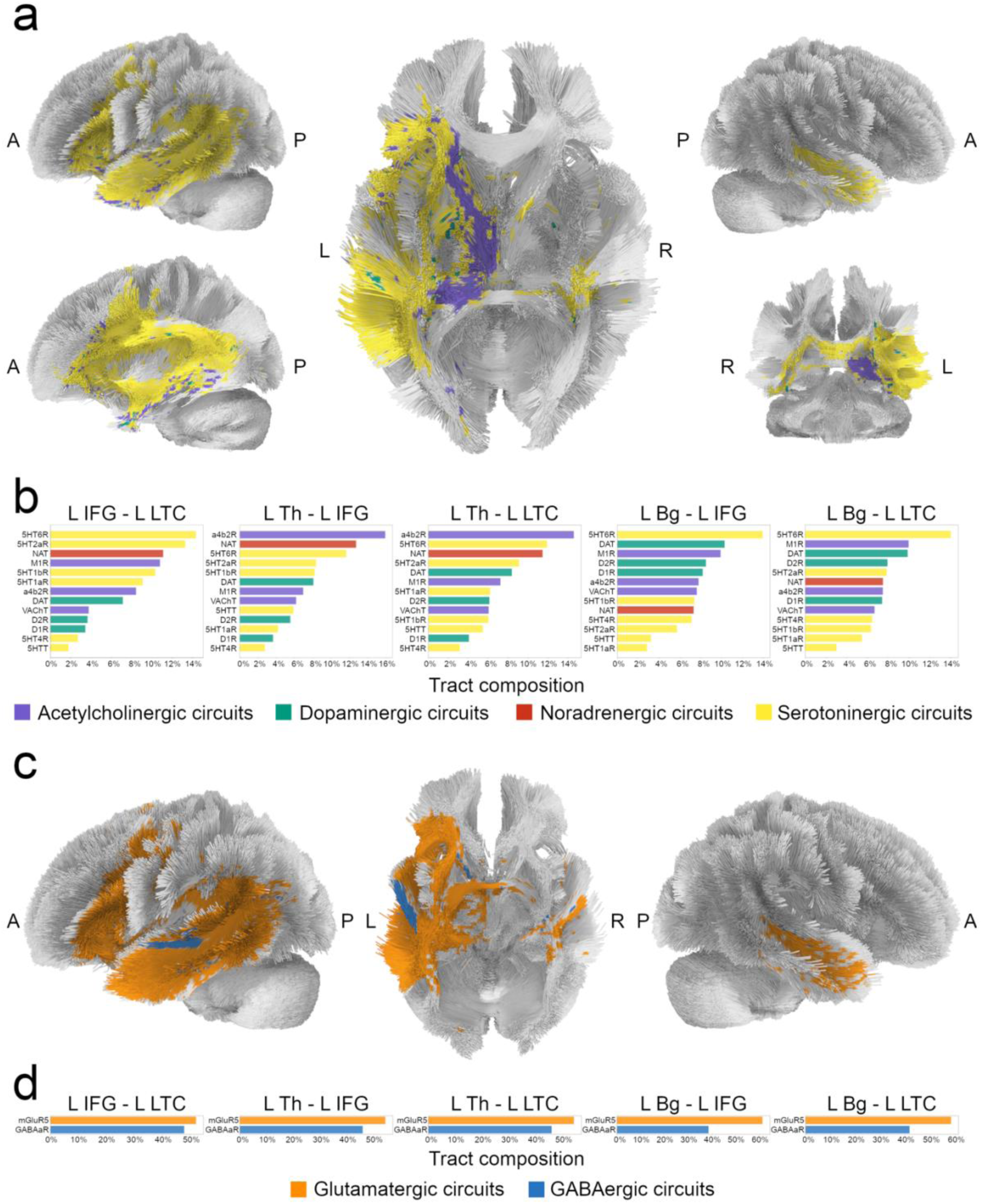
Neurotransmitter white matter projections of the language network. (a) Predominant modulatory neurotransmitter systems contributing to white-matter tracts connecting cortical and subcortical language clusters, based on neurotransmitter-weighted structural connectivity. Predominance reflects the neurotransmitter system with the highest relative contribution within each tract. (b) Relative receptor and transporter composition of selected language-related tracts for modulatory neurotransmitter systems. (c) Predominant fast-acting excitatory and inhibitory neurotransmitter systems (glutamate and GABA) across language-related tracts. (d) Relative glutamatergic and GABAergic composition of the same tracts. 5HT1aR, serotonin receptor 1a; 5HT1bR, serotonin receptor 1b; 5HT2aR, serotonin receptor 2a; 5HT4R, serotonin receptor 4; 5HT6R, serotonin receptor 6; 5HTT, serotonin transporter; A, anterior; a4b2R, acetylcholine receptor α4β2; D1R dopamine receptor 1, D2R dopamine receptor 2, DAT, dopamine transporter; L, left; L Bg, left basal ganglia; L IFG, left inferior frontal gyrus; L LTC, left lateral temporal cortex; L Th, left thalamus; M1R, muscarinic 1 receptor; NAT, noradrenaline transporter; P, posterior; R, right; R LTC, right lateral temporal cortex; VAChT, acetylcholine vesicular transporter.

The projection tracts from the left thalamus to both the left inferior frontal gyrus and the left lateral temporal cortex were predominantly cholinergic in contrast to cortico-cortical pathways (Figure 2a), especially rich in postsynaptic α4β2R cholinergic fibres (Figure 2b).

The projection tracts from the left basal ganglia (lentiform and caudate nuclei) to the left inferior frontal gyrus and the left lateral temporal cortex were also characterised by postsynaptic 5HT6R serotonergic fibres, followed by presynaptic dopaminergic and postsynaptic M1R cholinergic fibres (Figure 2b).

Regarding fast-acting excitatory and inhibitory neurotransmitters, glutamatergic fibres predominated overall relative to GABAergic fibres (Figure 2c-d). The reverse pattern was observed in a left anterior temporal region and in specific tracts of the anterior limb of the internal capsule (Figure 2c).

### 2.2. Neurotransmitter circuit disruption in post-stroke aphasia

#### 2.2.1. Patterns of disruption

To investigate patterns of neurotransmitter circuit disruption following stroke, we used the lesion data from the Aphasia Recovery Cohort.^65^ This dataset includes lesion masks from 196 adult patients who experienced aphasia at least six months after left-hemisphere ischemic stroke, and 31 patients without aphasia. Language deficits were assessed using the Western Aphasia Battery (WAB) or the Western Aphasia Battery-Revised (WAB-R).^66,67^

To calculate the impact of stroke lesion on neurotransmitter circuits, we used NeuroT-Map (https://github.com/Pedro-N-Alves/NeuroT-Map, version 2.0).^38^ This tool estimates the proportion of each receptor and transporter system affected by a focal lesion by overlaying lesion masks onto neurotransmitter location density and white matter atlases and calculates neurotransmitter injury ratios. These ratios are based on the neurobiological principle that receptors are predominantly postsynaptic, located on dendritic membranes of the postsynaptic neuron, whereas transporters are primarily presynaptic, located on axon terminal membranes of the presynaptic neuron. Accordingly, receptor location and tract maps serve as surrogates for postsynaptic neurons, while transporter maps represent presynaptic neurons within a given neurotransmitter system.^38^ The ratios thus reflect the relative extent of pre- and postsynaptic injury. A presynaptic ratio greater than 1 indicates more damage to presynaptic than postsynaptic structures in that circuit, whereas a postsynaptic ratio greater than 1 implies the opposite.^38^

Figure 3a represents the lesion overlap map, and Figure 3b the distribution of the maximum synaptic ratios (i.e., the most strongly disrupted neurotransmitter system per patient) of poststroke aphasia cases.

**Figure 3.**
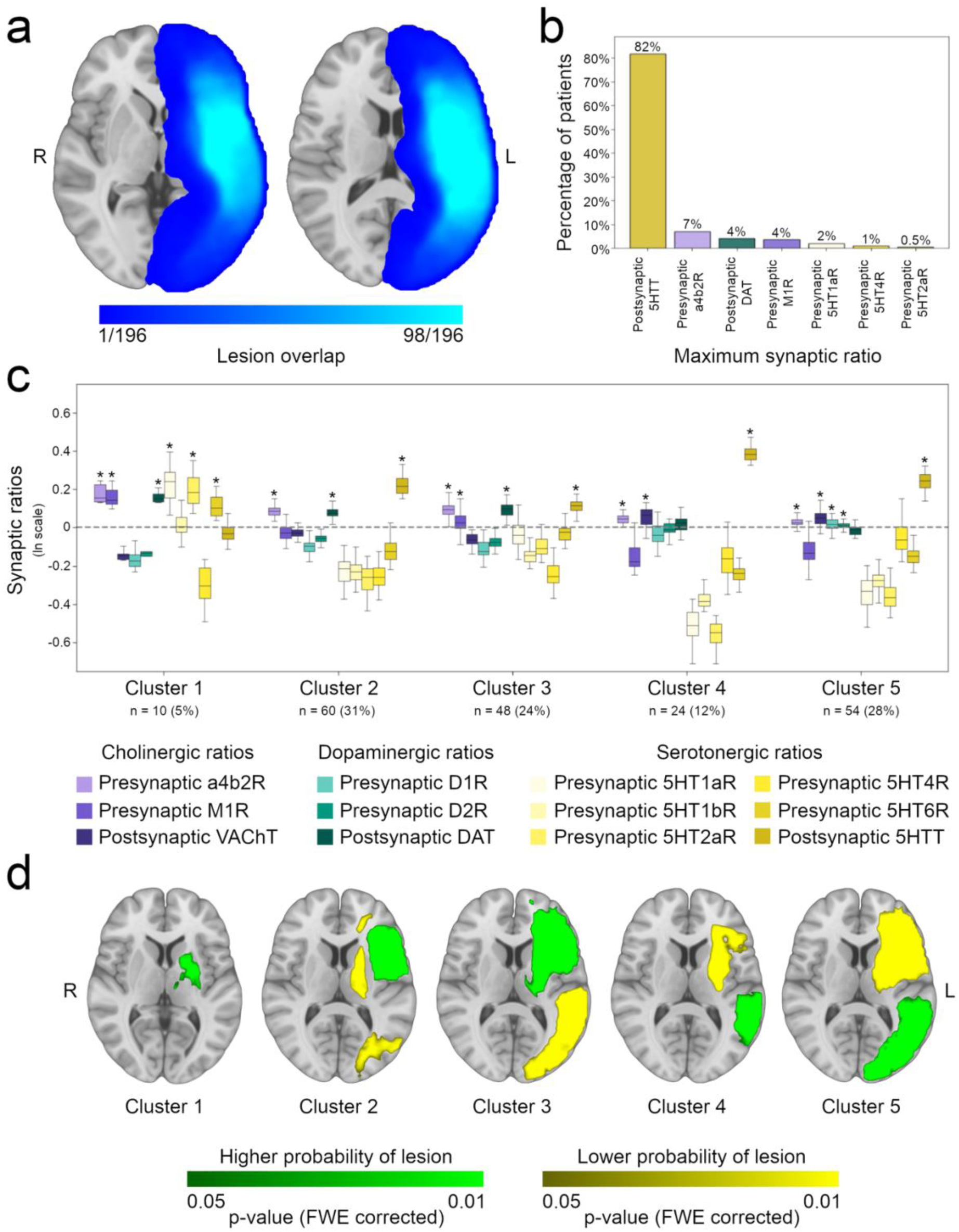
Poststroke aphasia neurochemical patterns. (a) Poststroke aphasia lesion overlap map (left, z=1; right, z=14). (b) Distribution of the maximum synaptic ratios. (c) Neurochemical clusters of poststroke aphasia; boxplots represent the distribution of synaptic ratios, in a natural logarithmic scale; the asterisks represent the distributions that are statistically significant positive (FDR-corrected p-values <0.05). (d) Statistical maps of the comparison between the lesion topography maps of each neurochemical cluster with the remaining clusters, controlling for lesion volume (cluster 1, z=5; clusters 2-5, z=9). 5HT1aR, serotonin receptor 1a; 5HT1bR, serotonin receptor 1b; 5HT2aR, serotonin receptor 2a; 5HT4R, serotonin receptor 4; 5HT6R, serotonin receptor 6; 5HTT, serotonin transporter; A, anterior; a4b2R, acetylcholine receptor α4β2; D1R dopamine receptor 1, D2R dopamine receptor 2, DAT, dopamine transporter; FWE, family-wise error; L, left; M1R, muscarinic 1 receptor; P, posterior; R, right; VAChT, acetylcholine vesicular transporter.

The distributions of percentage damage to receptor and transporter density maps, tract maps, and pre- and postsynaptic injury ratios in stroke patients with and without aphasia are presented in Supplementary Figures 2 and 3.

Patients with aphasia showed higher injury ratios across multiple neurotransmitter systems, including presynaptic cholinergic α4β2R, postsynaptic cholinergic VAChT, postsynaptic dopaminergic DAT, and postsynaptic serotonergic 5HTT compared with patients without aphasia. Conversely, lower injury ratios were observed in presynaptic cholinergic M1R, presynaptic dopaminergic D1R, and presynaptic serotonergic 5HT1a, 5HT1b, 5HT2a, and 5HT4 injury ratios (Benjamini–Hochberg False Discovery Rate (FDR)-corrected p-values<0.05).

The injury ratios of poststroke aphasia patients showed a tendency to be organized in clusters (Hopkins’ score = 0.13). The number of clusters was defined as 5, following the Elbow method. Figure 3c shows the K-means clustering of these ratios. Clustering was used here as a descriptive tool to summarise heterogeneity in neurochemical disruption profiles, rather than to define discrete biological subtypes.

Across clusters, variation was driven primarily by the balance between pre- and postsynaptic serotonergic and cholinergic disruption, with dopaminergic contributions modulating specific profiles. Cluster 1 differed from the other clusters in its serotonergic ratio pattern. While cluster 1 exhibited positive serotonergic presynaptic ratios—particularly for 5HT1aR, 5HT2aR, and 5HT6R—the remaining clusters showed predominantly postsynaptic serotonergic damage. Cluster 1 was also characterised by positive cholinergic presynaptic ratios involving α4β2R and M1R, as well as positive dopaminergic postsynaptic ratios. Clusters 2 and 3 both showed positive dopaminergic postsynaptic and cholinergic presynaptic ratios. In cluster 2, this involved only α4β2R, whereas in cluster 3 it included both α4β2R and M1R. Cluster 4 was characterised by a combination of positive cholinergic postsynaptic and presynaptic α4β2R ratios. In cluster 5, this combination co-occurred with positive dopaminergic presynaptic D1R and D2R ratios.

Anatomically, cluster 1 was characterised by lesions affecting the thalamus and lenticular nucleus. Cluster 2 was associated with fronto-insular damage, with relative sparing of the thalamic, parieto-occipital, and frontal paramedial regions. Cluster 3 comprised lesions involving the frontal cortex, thalamus, and basal ganglia, while parieto-occipital areas were largely preserved. Cluster 4 was defined by predominant temporal lobe damage, with sparing of the inferior frontal and basal ganglia regions. Finally, cluster 5 was characterised by temporo-occipital lesions, with relative preservation of frontal, thalamic, and basal ganglia structures.

The distribution of aphasia types across neurochemical clusters is shown in Table 1. Conduction aphasia was significantly associated with neurochemical cluster 4 (FDR-corrected p = 0.004). No other significant associations were observed.

**Table 1.**
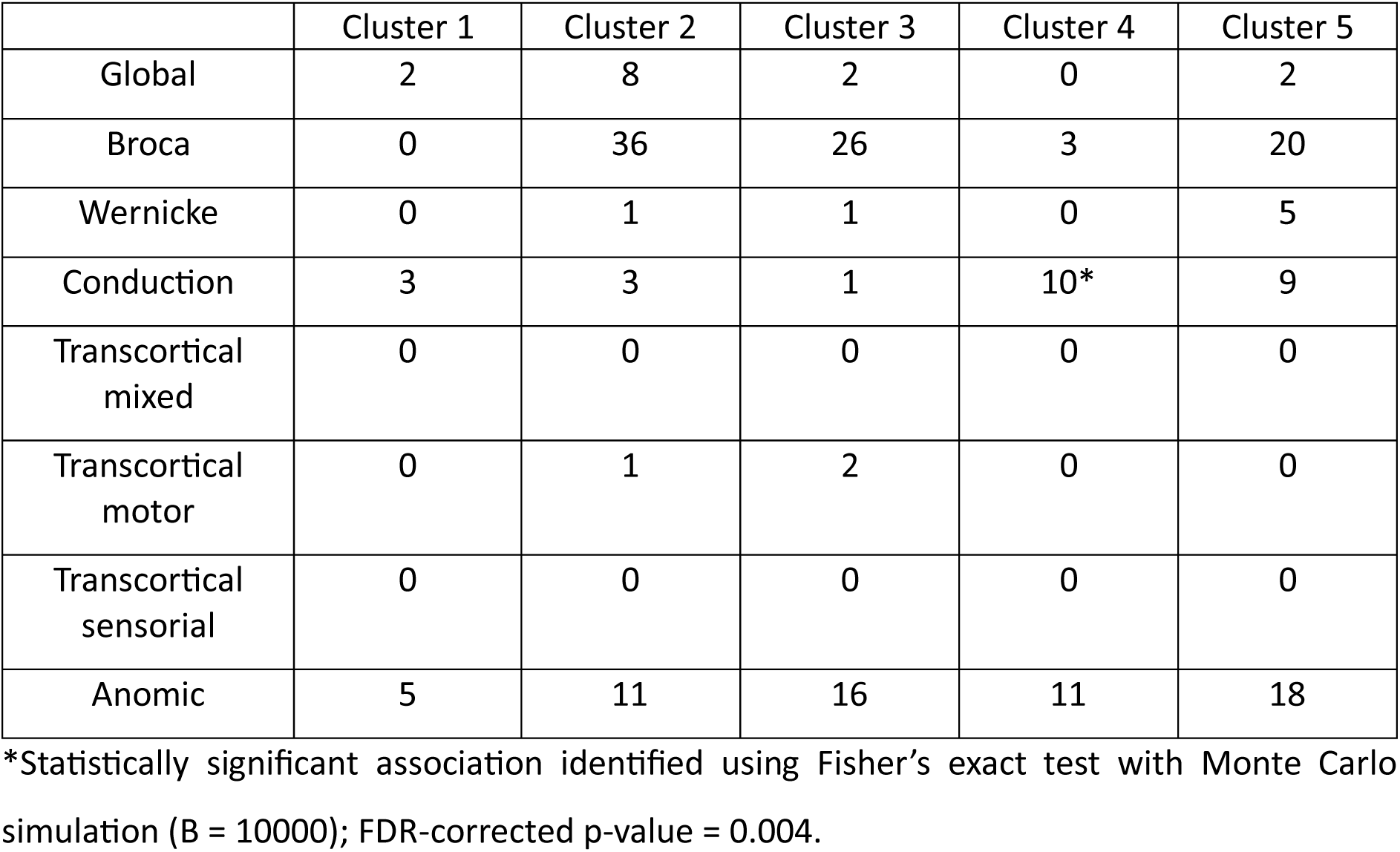
Distribution of aphasia types.

|  | Cluster 1 | Cluster 2 | Cluster 3 | Cluster 4 | Cluster 5 |
| --- | --- | --- | --- | --- | --- |
| Global | 2 | 8 | 2 | 0 | 2 |
| Broca | 0 | 36 | 26 | 3 | 20 |
| Wernicke | 0 | 1 | 1 | 0 | 5 |
| Conduction | 3 | 3 | 1 | 10* | 9 |
| Transcortical mixed | 0 | 0 | 0 | 0 | 0 |
| Transcortical motor | 0 | 1 | 2 | 0 | 0 |
| Transcortical sensorial | 0 | 0 | 0 | 0 | 0 |
| Anomic | 5 | 11 | 16 | 11 | 18 |

\*Statistically significant association identified using Fisher's exact test with Monte Carlo simulation (B = 10000); FDR-corrected p-value = 0.004.

#### 2.2.2. Neurochemical profiles of lobar and subcortical lesions

For the analysis of subcortical aphasia, we supplemented the Aphasia Recovery Cohort with 12 consecutive cases from a local hospital-based cohort. All had aphasia following ischemic or haemorrhagic stroke, available MRI, and lesions primarily involving the basal ganglia.

Lobar lesions in post-stroke aphasia (Figure 4a) demonstrated a neurochemical profile distinct from that of subcortical lesions (Figure 4b). In lobar lesions, serotonergic postsynaptic injury predominated (Figure 4c). In contrast, subcortical lesions showed predominant injury in presynaptic receptors, specifically serotonergic 5-HT1aR and 5-HT2aR, and cholinergic α4β2 and M1 subtypes (Figure 4d).

**Figure 4.**
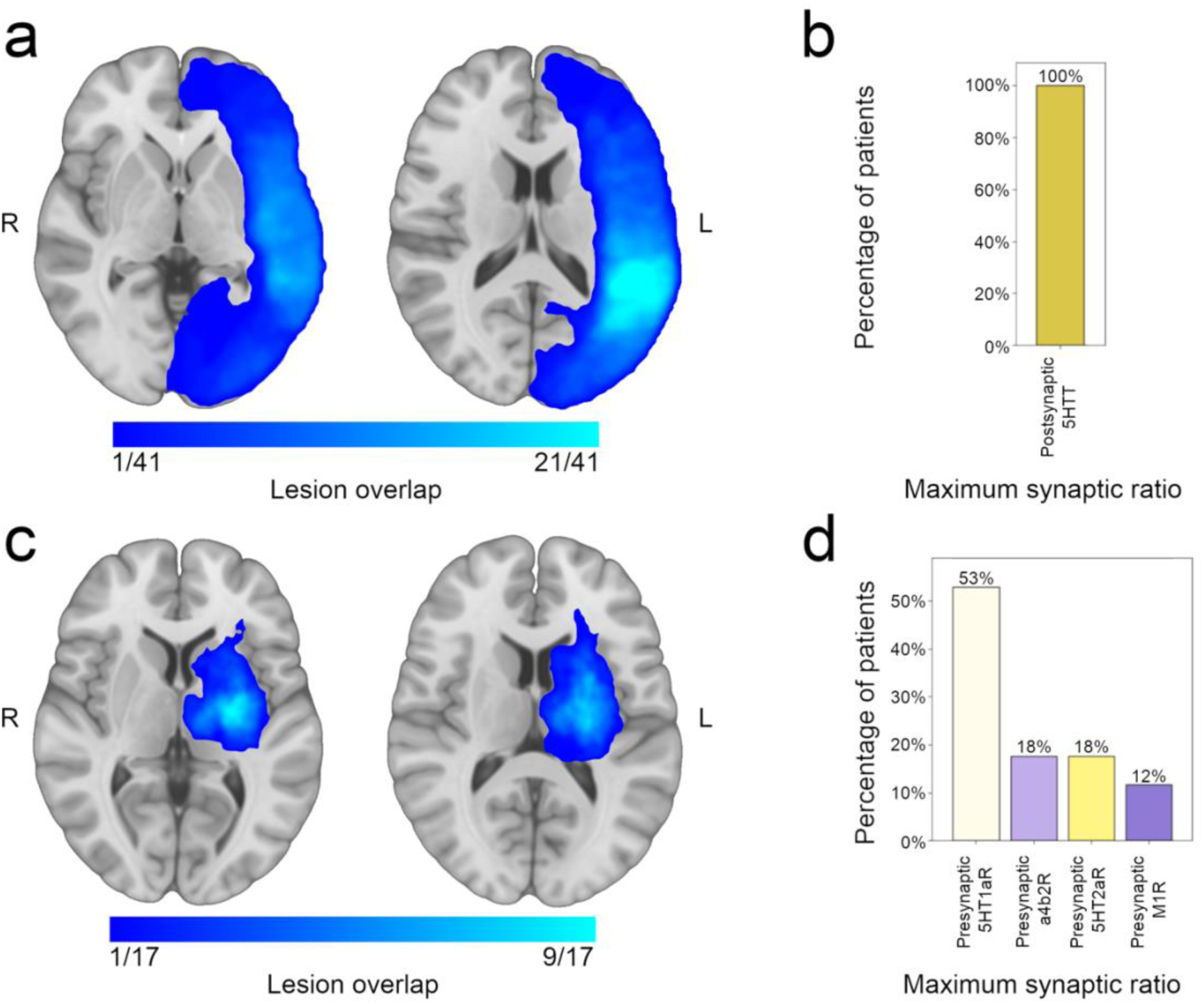
Neurochemical profiles of lobar and subcortical poststroke aphasia lesions. (a) Lesion overlap map of lobar lesions. (b) Distribution of the maximum synaptic ratios of lobar lesions. (c) Lesion overlap map of subcortical lesions. (d) Distribution of the maximum synaptic ratios of subcortical lesions. 5HT1aR, serotonin receptor 1a; 5HT2aR, serotonin receptor 2a; 5HTT, serotonin transporter; a4b2R, acetylcholine receptor α4β2; L, left; M1R, muscarinic 1 receptor; R, right.

#### 2.2.3. Association between neurochemical measures and language outcomes

We also investigated how disruptions in neurotransmitter circuits and neurochemical clusters were associated with WAB scores (0-100; higher is better). The strongest regression coefficients in the regularised model indicate that greater damage to the cholinergic presynaptic system was associated with lower fluency and spontaneous speech scores, whereas greater damage to the serotonergic postsynaptic 5HT1aR system was associated with lower sentence completion scores (Figure 5a).

**Figure 5.**
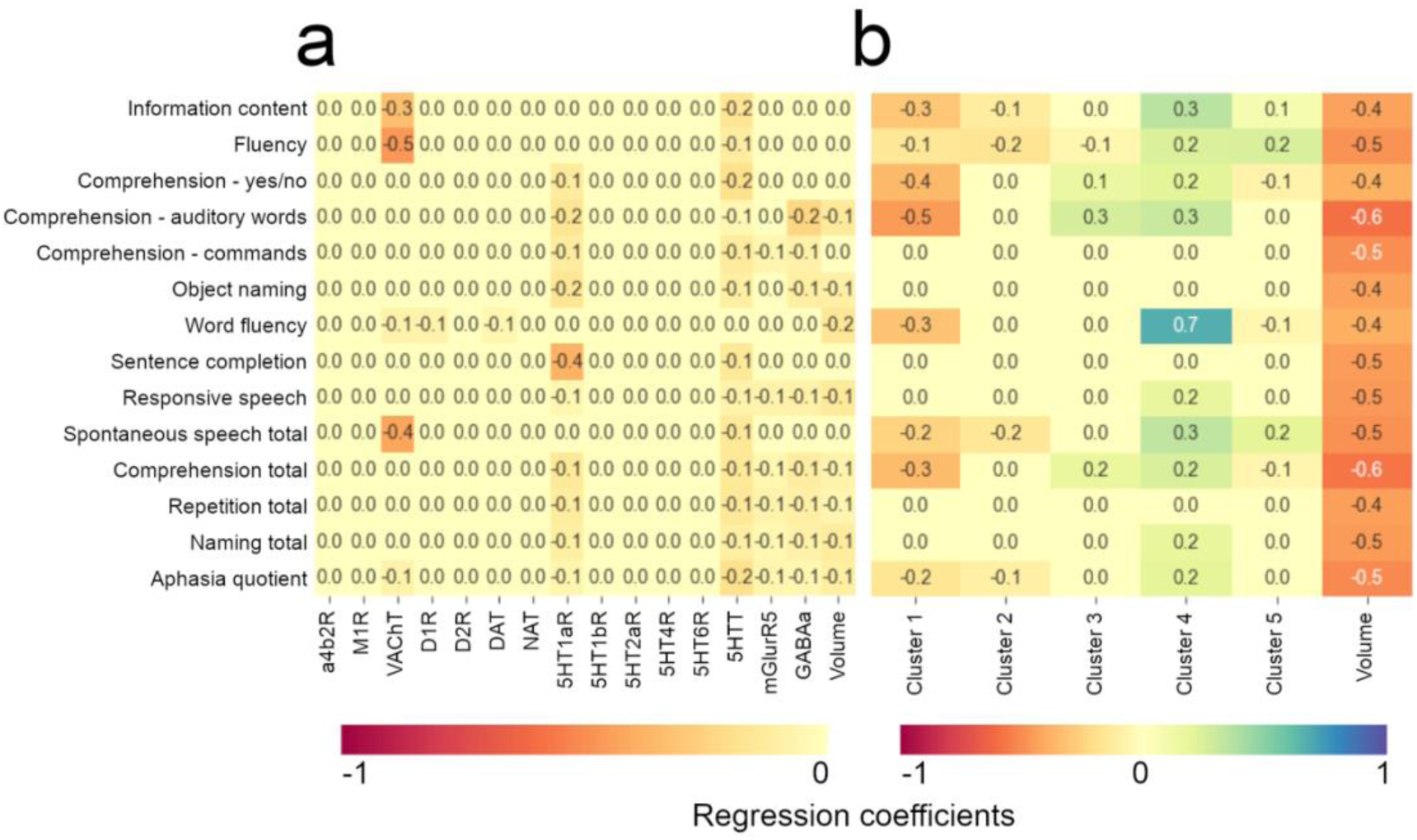
Elastic Net regularised regression between neurochemical measures (predictors) and language outcomes. (a) Predictors included the volume of damage to each neurotransmitter system and the total lesion volume. (b) Predictors included the neurochemical clusters and the total lesion volume. Numbers in the matrices represent regression coefficients. 5HT1aR, serotonin receptor 1a; 5HT1bR, serotonin receptor 1b; 5HT2aR, serotonin receptor 2a; 5HT4R, serotonin receptor 4; 5HT6R, serotonin receptor 6; 5HTT, serotonin transporter; A, anterior; a4b2R, acetylcholine receptor α4β2; D1R dopamine receptor 1, D2R dopamine receptor 2, DAT, dopamine transporter; L, left; M1R, muscarinic 1 receptor; NAT, noradrenaline transporter; P, posterior; R, right; VAChT, acetylcholine vesicular transporter.

When considering neurochemical clusters as predictors, cluster 4 was associated with better fluency, comprehension, and naming scores, whereas cluster 1 was associated with worse fluency and comprehension (Figure 5b).

### 2.3. *Post hoc* analysis of clinical trials

To investigate the hypothesis that neurotransmitter ratios generated by NeuroT-Map can predict pharmacological responses in patients with post-stroke aphasia, we conducted a *post hoc* analysis of clinical trials assessing the efficacy of acetylcholine, dopamine, or serotonin modulators in several aphasic populations.

We collected language assessment and lesion data from 56 patients with post-stroke aphasia across four clinical trials. In 36 patients, the outcome measure was the WAB Aphasia Quotient (AQ) severity score: 30 patients received the acetylcholinesterase inhibitor donepezil across two trials, and 6 received the selective serotonin reuptake inhibitor sertraline.^68–71^ In the remaining 20 patients, the outcome measure was the Comprehensive Aphasia Test comprehension score, and all were treated with donepezil.^50,72^

We then performed a *post hoc* analysis using data from the four trials. The predictive variable was the match between the drug used in each trial and the highest neurotransmitter ratio generated by NeuroT-Map. Specifically, matching for donepezil was based on presynaptic ratios of the acetylcholine circuit (either α4β2 or M1 systems), while matching for sertraline was based on the postsynaptic ratio of the serotonergic system. Regarding outcome measures, for the WAB-AQ we defined improvement as an increase of at least 10 points, which corresponds to the intermediate value between the thresholds used in the original trials (5 and 15).^69,71^ Although there is no consensus on a definitive cut-off, a 10-point increase is considered a robust marker of clinical improvement.^73–75^ For the CAT comprehension score, any positive change was considered an improvement, consistent with the original study.^50^ We observed that in this hypothesis-generating, exploratory analysis, patients were more likely to improve when the administered drug matched the predominant neurotransmitter ratio identified by NeuroT-Map (Table 2; χ² = 4.129, p = 0.042; Cramer’s V = 0.272; moderate effect size).

**Table 2.**
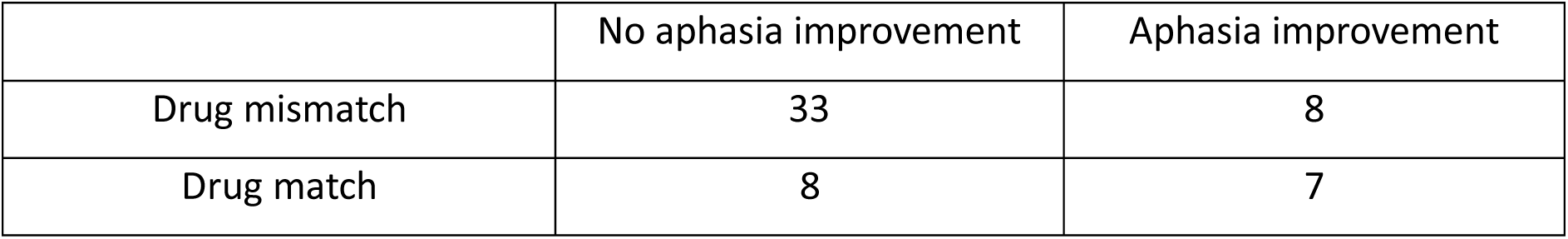
Contingency table showing the relationship between drug–neurotransmitter matching (based on the maximum neurotransmitter ratio identified by NeuroT-Map) and aphasia improvement.

A more conservative analysis using stricter thresholds for defining aphasia improvement was also performed: 15 points for the WAB-AQ and 5 points for the CAT comprehension score. It also revealed a significant association (Supplementary Table 1; Fisher’s exact test, p = 0.026; Cramer’s V = 0.329; moderate effect size). These analyses are intended to generate mechanistic hypotheses rather than to establish clinical efficacy.

## 3. Discussion

In this study, we characterised the neurotransmitter circuits of the human language network and examined how their disruption relates to post-stroke aphasia and treatment response. By integrating large-scale functional mapping with neurotransmitter-informed structural connectivity, we show that language circuits exhibit a non-uniform neurochemical organisation: cortico–cortical pathways are preferentially enriched in serotonergic and glutamatergic signalling, thalamo–cortical projections are predominantly cholinergic, and anterior temporal connections show relative enrichment of GABAergic circuits. Extending this framework to focal brain injury, we observed that aphasia following lobar stroke is most commonly associated with postsynaptic serotonergic disruption, whereas subcortical aphasia shows a distinct profile characterised by presynaptic serotonergic or cholinergic involvement. Finally, in a *post hoc* analysis of pharmacological trials, improvement was more frequent when treatment matched the dominant neurotransmitter imbalance inferred from the lesion profile. Together, these findings suggest that the organisation and disruption of language networks can be framed in neurochemical terms, providing a conceptual bridge between circuit disconnection, neurotransmitter imbalance, and heterogeneity in treatment response after stroke.

An important consideration in interpreting these findings is that our neurochemical measures are derived from normative, PET-based maps of receptor and transporter distributions,^37^ projected onto structural connectivity priors,^63^ rather than from participant-specific neurotransmitter imaging. As such, our approach does not quantify absolute neurotransmitter concentrations in individual patients but instead estimates the relative vulnerability of neurotransmitter-specific circuits to focal lesions. This framework rests on the reproducibility of large-scale receptor and transporter topographies across individuals and on the principle that lesions interrupt network elements in proportion to their underlying neurochemical composition.^76^ In this sense, neurotransmitter-weighted disconnection provides a biologically informed proxy for the likelihood that a given lesion disproportionately affects pre- or postsynaptic components of specific neuromodulatory systems. Future work should quantify sensitivity to alternative connectome priors, tractography algorithms ^77,78^, receptor map sources, and genetics^79^, and should test the extent to which individual variability (e.g., age, sex and vascular risk factors) modulates neurotransmitter-weighted vulnerability estimates. Our results are consistent with receptor autoradiography observations showing that language areas cluster in receptor expression patterns, despite their cytoarchitectonic and topographic diversity.^34^ They further reported that the main discriminants between language and non-language areas were the serotonergic 5HT1aR and cholinergic muscarinic receptor densities. In our analysis, serotonergic and glutamatergic circuits predominated across most language network connections, with two notable exceptions: thalamo-cortical circuits, which were mainly cholinergic, and anterior temporal lobe fibres, which showed a GABAergic predominance. This pattern contrasts with the broader diversity of predominant neurotransmitter circuits across the brain previously described using the same method.^38^ The predominance of GABAergic fibres in the left anterior temporal regions may help explain the association between GABA concentrations in anterior temporal areas, as measured by MR spectroscopy, and semantic processing.^35,36^

The characterisation of patterns of neurotransmitter circuit disruption sheds light on the neurochemical basis of poststroke aphasia and opens new opportunities for neuropharmacological modulation. In most patients, we observed a predominant imbalance of the serotonergic system, with preferential disruption of postsynaptic circuits. Supporting this view, Hillis and colleagues reported two observational studies showing that the use of selective-serotonin reuptake inhibitors was strongly associated with improved recovery of naming in patients with poststroke aphasia.^58–61^ These findings are consistent with our neuropharmacological hypothesis. Selective serotonin reuptake inhibitors act on presynaptic serotonin transporters, which, according to our results, are relatively more frequently spared compared to postsynaptic structures. The negative results of large clinical trials are likely due to not pairing the investigative drugs with adequate doses of behavioural therapies. All of the large-scale drug trials of patients with stroke have been pragmatic, meaning that the patients only received standard levels of therapist-delivered interventions, which, in most health-care systems, are not enough to ‘move the dial’ on motor or language outcomes.^58–60^ The cluster analysis showed that serotonergic postsynaptic deficits may be accompanied by distinct patterns of pre- or postsynaptic cholinergic and dopaminergic imbalances, suggesting the existence of different neurochemical fingerprints across this group of patients.

Importantly, patients with subcortical lesions showed a strikingly different neurochemical profile, characterised by predominant presynaptic disruptions of serotonergic or cholinergic circuits. Acetylcholinesterase inhibitors, which increase acetylcholine levels in the synaptic cleft, have been associated with improvements in language outcomes.^49–52^ According to our neuropharmacological model, this effect may be driven by subcortical lesions, as this class of drugs would be particularly effective in the context of presynaptic-predominant disruptions. This hypothesis is consistent with the observations of Hong and colleagues, who conducted a randomised controlled trial of galantamine in chronic poststroke aphasia.^49^ Galantamine was associated with improvements in aphasia quotient, with subcortical-dominant lesion patterns emerging as a determinant of good responsiveness. To our knowledge, there are no data on the use of drugs acting on postsynaptic serotonergic circuits in patients with post-stroke aphasia.^49^

The *post hoc* analysis of clinical trials using neurotransmitter modulator drugs in patients with post-stroke aphasia supports the hypothesis that serotonin selective reuptake inhibitors may have higher efficacy in patients with predominant serotonergic postsynaptic injury imbalance and acetylcholinesterase inhibitors in patients with predominant presynaptic cholinergic injury imbalance. These preliminary findings provide proof of concept that neurotransmitter-informed lesion profiling may help guide drug selection in post-stroke aphasia and motivate prospective, stratified trials to rigorously test this hypothesis.

An important caveat is that the mechanism by which neurotransmitter-to-drug matching would translate into differential clinical outcomes remains to be established. Our model proposes that when a lesion disproportionately destroys presynaptic terminals, as in subcortical lesions with cholinergic predominance, postsynaptic receptors remain relatively intact, and drugs that increase synaptic neurotransmitter availability (e.g., acetylcholinesterase inhibitors) may then drive activity through these preserved targets. In contrast, when postsynaptic structures are preferentially damaged, presynaptic reuptake inhibitors may maintain higher synaptic concentrations, thereby recruiting partially spared downstream receptors. Critically, however, these pharmacological effects are unlikely to operate in isolation. Neuromodulatory systems are thought to lower the threshold for synaptic plasticity and learning, and their therapeutic effects are therefore expected to be amplified by concurrent behavioural practice at sufficient intensity. The trials included in the present *post hoc* analysis varied considerably in the dose and timing of speech and language therapy delivered alongside pharmacological treatment, an important source of response heterogeneity independent of neurochemical profiling. Future stratified trials should explicitly pair neurochemical lesion classification with controlled, adequate doses of therapist-delivered intervention in order to properly test this mechanistic hypothesis.

Beyond *post hoc* analyses, the present framework naturally lends itself to prospective testing in stratified clinical trials. A straightforward design would involve classifying patients based on their dominant neurotransmitter disruption profile—such as serotonergic versus cholinergic predominance, or pre- versus postsynaptic imbalance—prior to randomisation, using standard structural MRI and lesion mapping. Pharmacological interventions could then be targeted to these profiles, with outcome measures focusing on domain-specific language functions rather than global communication scales alone. Such enrichment strategies may reduce response heterogeneity and increase statistical power, offering a principled alternative to one-size-fits-all pharmacological approaches in post-stroke aphasia.

The association between neurochemical clusters and aphasia classification derived from the WAB was limited, and neurochemical measures showed weak predictive value for performance across individual language domains. This result was expected, as previous evidence indicates that neurotransmitter systems have multifaceted roles and are not directly tied to any single behaviour or cognitive function.^37,38^ In the context of degenerative disorders of language, for instance, grey matter atrophy has been shown to spatially co-localise with serotonin, dopamine, and glutamatergic pathways.^80^ In line with our results, similar neurotransmitter patterns have been observed across the agrammatic and semantic variants of primary progressive aphasia, despite their distinct language dysfunction profiles.^80^ Classifying aphasia in the chronic phase may have also contributed to weaker associations, as recovery patterns are more heterogeneous at this stage.^22^ In our data, a single statistically robust association emerged: neurochemical cluster 4 was associated with conduction aphasia.

Notably, this cluster was also characterised by relatively better fluency and comprehension scores, proving converging support for its behavioural relevance.

Despite these advances, a number of limitations warrant consideration. First, the *post hoc* analyses were constrained by data availability. Some of the included clinical trials were conducted many years ago, precluding access to lesion data for all participants. In addition, the trials differed in outcome measures, were predominantly conducted in chronic patients, and in some cases lacked placebo controls. While these factors may limit causal inference, the present results nevertheless provide a principled framework for the design of future, prospectively stratified trials. Second, aphasia characterisation relied on the WAB. Although the WAB is a widely used and well-validated instrument for diagnosing and classifying post-stroke aphasia, it may be too conservative and can underestimate residual language impairment and primarily reflects syndromic categorisation rather than dissociable psycholinguistic components such as phonology, lexical access, and semantics.^81^ Third, our study does not assess the impact of language lateralisation. Language regions in participants with right-hemisphere lateralisation are underrepresented in the group-level analysis, as the lesion study included only patients with left-hemisphere strokes. Finally, functional delineation of the language network was derived from a localiser contrasting sentence reading with nonword reading.^82,83^ While this approach is widely published, it does not dissociate distinct psycholinguistic operations within the language system, which may engage partially overlapping but neurochemically distinct circuits. In particular, the localiser approach was designed to robustly identify domain-general language-responsive cortex rather than to isolate specific psycholinguistic operations. As such, the present analyses do not distinguish between processes such as phonological encoding, lexical access, or semantic integration, which may rely on partially overlapping but neurochemically distinct circuits. Regions such as the insula did not reach the predefined prevalence threshold across individuals and were therefore not included in the analysis. Future work combining neurotransmitter-informed circuit mapping with task paradigms explicitly targeting these operations will be important to refine the functional–neurochemical specificity of language networks.

In conclusion, this study delineates the neurochemical architecture of the human language network and demonstrates that post-stroke aphasia is associated with distinct patterns of neurotransmitter circuit disruption that vary with lesion topology. By linking serotonergic, cholinergic and dopaminergic imbalances to structural disconnection and behavioural outcomes, our results provide a mechanistic framework that bridges systems neuroscience and clinical neuropharmacology. Although exploratory, the convergence between neurotransmitter-specific lesion profiles and pharmacological response across independent trials suggests that circuit-level neurochemical mapping may offer a principled route toward stratifying patients and designing targeted intervention studies in post-stroke aphasia.

## 4. Methods

### 4.1. Neurotransmitter circuits of language network

#### 4.1.1. Cortical regions of the language network

We used an *f*MRI dataset comprising 806 healthy participants (93% right-handed, 5% left-handed, 2% ambidextrous) performing language tasks, provided by Lipkin and colleagues, to define the cortical regions of the language network.^62^ The majority (n = 688) performed a sentence-reading task contrasted with nonword reading. The remaining participants completed one of several tasks: a reading task contrasting sentences, wordlists, and nonwords (n = 82); a reading task contrasting sentences, wordlists, nonwords, and Jabberwocky (n = 23); a listening task contrasting intact with degraded speech (n = 8); a listening task contrasting sentences, wordlists, nonwords, and Jabberwocky (n = 4); or a listening task contrasting sentences with nonwords (n = 1).^62^ Despite this variation, these paradigms were shown to produce broadly similar activation maps.^62^

Individual t-maps were thresholded at p<0.05. To identify consistent language regions across individuals, we used the FSL tool “cluster”, applying a threshold of 0.5 (i.e., voxels were included if they were part of the language map in at least 50% of individuals) and a minimum cluster extent of 10 voxels (2×2×2 mm resolution).

#### 4.1.2. Subcortical regions of the language network

Functional inter-individual alignment was performed using the Advanced Normalization Tools (ANTs) script “buildtemplateparallel.sh”.^23,45,84^ Four iterative diffeomorphic transformations were estimated to align individual language-task t-maps to a common template.^84^ Cross-correlation was used as the similarity metric, and the Greedy SyN algorithm was employed for the transformation. The resulting warp fields were applied to the MNI152-aligned language maps using the ANTs script “WarpImageMultiTransform”, enabling the representation of all 806 individual language maps in the common functional space. This functional template was derived after initial MNI152 anatomical registration, ensuring correspondence with standard space while improving functional alignment.

As in the cortical analysis, individual t-maps were thresholded at p<0.05, and the FSL “cluster” tool was used to identify subcortical language regions. A minimum cluster extent of 10 voxels (2×2×2 mm resolution) was applied. To account for the greater inter-individual variability not fully resolved by functional alignment, a more permissive threshold of 0.25 was applied (i.e., voxels were included if they were present in at least 25% of individual maps).

The resulting language cluster map was visualized using Connectome Workbench 2.1.0 (https://www.humanconnectome.org/software/connectome-workbench).

#### 2.1.3 Projection of neurotransmitter receptor and transporter densities onto the language structural network

We used the Functionnectome to project neurotransmitter receptor and transporter densities from grey matter onto white matter based on voxel-wise structural connectivity probabilities.^38,63^

Normative structural probability maps, derived from whole-brain deterministic tractography of 7T diffusion-weighted MRIs in 176 healthy Human Connectome Project participants, were used to define structural connectivity priors.^64^ We included only those tracts linking two or more language clusters. The MRtrix commands “tck2connectome” and “connectome2tck” were used for tract selection.^85^ Input nodes consisted of language clusters that were dilated using the “maskfilter” command with the dilate option (2 iterations), which expands a binary mask by one voxel layer per iteration.^85^ This step ensured that the clusters intersected the grey–white matter interface.^86^ The resulting tractograms, linking specific pairs of language clusters, were then used as priors in the Functionnectome.

PET-derived density maps were used as inputs to the Functionnectome for the following neurotransmitter systems: acetylcholine receptors (α4β2R, M1R), acetylcholine transporter (VAChT), dopamine receptors (D1R, D2R), dopamine transporter (DAT), noradrenaline transporter (NAT), serotonin receptors (5HT1aR, 5HT1bR, 5HT2aR, 5HT4R, 5HT6R), serotonin transporter (5HTT), GABA receptor (GABAaR), and glutamate receptor (mGluR5).^37^ To account for variability in PET signal across tracers, each map was normalized to a 0–1 range, based on its own minimum and maximum values.^38^ This normalisation preserves relative spatial distributions within each tracer while avoiding scale differences across radioligands. When multiple tracer maps existed for the same receptor or transporter, their voxel-wise median was computed.^38^

In the resulting neurotransmitter-weighted structural maps, each white matter voxel value represents the average receptor or transporter density of the structurally connected grey matter voxels, weighted by the probability of connectivity. The relative contribution of each neurotransmitter system to a specific pairwise connection was quantified by dividing the sum of that system’s voxel values by the total voxel sum across the map.

DSI Studio was used to visualize the predominant neurotransmitter system within the language structural network.^87^

### 4.2. Patterns of neurotransmitter circuit disruption in post-stroke aphasia

#### 4.2.1. Stroke lesion datasets

We investigated post-stroke disruptions in neurotransmitter circuits using the Aphasia Recovery Cohort.^65^ This cohort includes lesion masks from a total of 227 adult patients: 196 had persistent aphasia at least six months after experiencing a left-hemisphere ischemic stroke (age, median [interquartile range]: 58 [50–66] years; sex: 72 female/124 male; time since stroke, median [interquartile range]: 703 [398–1611] days), and 31 did not have aphasia (age, median [interquartile range]: 59 [53–65] years; sex: 15 female/16 male; time since stroke, median [interquartile range]: 1076 [416–2660] days). Lesions were manually segmented on T1-weighted images and spatially normalised to the MNI152 standard space using the Clinical Toolbox for SPM.^88^

The WAB^67^ or WAB-R^66^, standard clinical instruments in aphasia evaluation, were used to assess language deficits through their four core subtests: a) spontaneous speech, rated based on information content and fluency; b) auditory verbal comprehension, including yes/no questions, auditory word recognition, and sequential commands; c) repetition, involving single words and phrases; d) naming and word finding, assessed through object naming, word fluency, sentence completion, and responsive speech. Higher scores on each subtest indicate better performance.^67^ Based on these scores, an aphasia type can be classified, and the AQ can be calculated, ranging from 0 (very severe aphasia) to 100 (no aphasia).^67^ The median [interquartile range] AQ score was 66 [42–91] in patients with aphasia and 98 [95–98] in patients without aphasia. All participants provided informed consent, and the study was approved by the Institutional Review Board of the University of South Carolina – Columbia.

To complement this dataset for the analysis of subcortical aphasia, we included cases from a local hospital-based cohort. We screened discharge notes from patients released from the Stroke Unit, Department of Neurology, Hospital de Santa Maria, Lisbon, between September 2019 and January 2023. The local cohort study was approved by the Centro Académico de Medicina de Lisboa Ethics Committee and conducted in accordance with local regulations. Patients with ischemic or haemorrhagic strokes, with MRI available, and whose lesions were centred in the basal ganglia, were selected (n = 12). Lesions were delineated on diffusion-weighted images (DWI) acquired on a Phillips Achieva 3.0 T or Philips Intera 1.5T scanner with a b-value of 1000 s/mm². To align the lesions to the MNI152 standard space, native-space DWI images were first linearly registered to the corresponding native T1-weighted images using FSL’s “flirt” command (12 degrees of freedom, mutual information cost function).^89^ Then, T1-weighted images were linearly and nonlinearly registered to MNI space using flirt (12 degrees of freedom, correlation ratio cost function) and “fnirt” (maximum of 5 non-linear iterations per resolution level).^89^

#### 4.2.2. Neurotransmitter circuit disruption analysis

We quantified the effects of focal stroke lesions on neurotransmitter systems using NeuroT-Map (version 2.0; https://github.com/Pedro-N-Alves/NeuroT-Map).^38^ This approach involves superimposing individual lesion masks onto atlases of neurotransmitter receptor and transporter density, as well as white matter maps. By computing the spatial overlap, the software estimates the percentage of each neurotransmitter circuit that is compromised by the lesion.^37^ Three outputs: a) the proportion of receptor and transporter location maps affected; b) the proportion of receptor and transporter tract maps affected; c) the pre- and postsynaptic injury ratios, presented on a logarithmic scale.

This analytical framework is based on converging neurobiological evidence indicating that neurotransmitter receptors are predominantly postsynaptic, whereas transporters are chiefly presynaptic. Consequently, receptor density maps serve as proxies for postsynaptic elements, and transporter maps reflect presynaptic components within each neurotransmitter system.

From these mappings, we derived ratios that reflect the comparative degree of pre- and postsynaptic disruption. A presynaptic ratio greater than 1 indicates that presynaptic structures are more extensively affected than their postsynaptic counterparts in a given circuit. Conversely, a postsynaptic ratio exceeding 1 reflects relatively greater involvement of postsynaptic elements.^38^

#### 4.2.3. Cluster analysis of the synaptic ratios

We have previously shown that synaptic ratios in stroke patients exhibit a strong clustering tendency.^38^ Here, we investigated whether the synaptic ratios of individuals with post-stroke aphasia follow a uniform distribution or similarly tend to form clusters.

First, we assessed the clustering tendency of the data using the Hopkins statistic, as implemented in the pyclustertend toolkit (https://pyclustertend.readthedocs.io/, version 1.8.2). Values close to 0 indicate a high tendency to form clusters, whereas values above 0.3 suggest a more uniform distribution. We then applied the “elbow” method, available in the Yellowbrick library (https://www.scikit-yb.org/, version 1.5), to estimate the optimal number of clusters, and performed unsupervised clustering using the k-means algorithm from the scikit-learn package (https://scikit-learn.org/, version 1.2.2).^90^ K-means clustering was chosen for its robustness and interpretability in low-dimensional ratio spaces. The resulting neurotransmitter ratio profiles for each cluster were visualized on a natural logarithmic scale.

To assess whether the distribution of aphasia types differed across the neurochemical clusters, we applied Fisher’s exact test using R, with Monte Carlo simulation (B = 10000) to approximate the p-value.^91^ Statistical significance was set at p<0.05. To identify which individual cells contributed most to the overall association, we computed standardized residuals from a chi-square test of independence. To control for multiple comparisons across all table cells, we applied the FDR correction using the Statsmodels library (https://www.statsmodels.org/, version 0.14.4).

To examine the association between lesion topography and neurochemical cluster membership, voxel wise non-parametric permutation testing was performed using FSL randomise (5000 permutations) within a general linear model framework. An omnibus F-contrast was specified to test for between-cluster effects.^92^ Threshold-free cluster enhancement was applied, and statistical significance was determined using family-wise error correction at p < 0.05. In regions demonstrating significant omnibus effects, post hoc voxel wise t-contrasts were conducted to compare each cluster against the remaining clusters, using identical permutation and correction parameters.

#### 4.2.4. Association between neurochemical measures and language outcomes

Multicollinearity was present among the neurotransmitter measures, as larger lesions tend to disrupt multiple circuits simultaneously and neurotransmitter circuits show spatial correlations.^38^

To examine the independent contributions of neurotransmitter measures to WAB outcomes while addressing multicollinearity and enabling variable selection, we used Elastic Net regularized regression with internal cross-validation (https://scikit-learn.org/, version 1.2.2).^90^ The regularization parameters were optimized using 5-fold cross-validation (cv=5), exploring a grid of α values (alphas=np.logspace(-3, 2, 100)) and L1 ratios (l1_ratio=[0.1, 0.3, 0.5]). Higher L1 ratios were avoided due to their reduced stability under multicollinearity. The model was fit with a maximum of 50.000 iterations (max_iter=50000) and a convergence tolerance of 1e-5 (tol=1e-5). Although this approach does not yield p-values, it provides regularized coefficient estimates optimized through cross-validation, enabling stable variable selection and reducing coefficient instability that arises from multicollinearity among predictors.

We ran two models. In the first, predictors were the proportions of each neurotransmitter circuit disrupted by the lesion, based on the maximum disruption values from location and tract maps. In the second, predictors were the patient’s neurochemical cluster assignments. All continuous predictors were standardised using StandardScaler from sklearn.preprocessing, while the binary cluster variables were left unscaled (https://scikit-learn.org/, version 1.2.2). In the first model, we constrained coefficients to be non-positive (positive=True with predictor sign flipping) to reflect biological plausibility—greater disruption of neurotransmitter circuits is unlikely to improve language outcomes—and to prevent spurious sign reversals driven by shared variance among predictors.^93,94^ Lesion volume was included in both due to its impact in stroke neuroimaging measures.^95^

Data visualization was performed with Matplotlib (https://matplotlib.org/, version 3.7.1) and Seaborn (https://seaborn.pydata.org/, version 0.11.1).

### 4.3. *Post hoc* analysis of post-stroke aphasia clinical trials

We performed a *post hoc* analysis of clinical trials evaluating acetylcholine, dopamine, or serotonin modulators in patients with post-stroke aphasia to test whether neurotransmitter ratios derived from NeuroT-Map could predict pharmacological responses.

Following a literature search and data requests, we obtained lesion maps and language outcomes from four clinical trials. In three of these, the WAB-AQ was the primary outcome measure; in one, the primary outcome was the comprehension score of the CAT.

One of the trials by Berthier and colleagues was an open-label study evaluating the use of donepezil and intensive language action therapy in 10 patients with mild to moderate aphasia of more than six months’ duration.^69^ The intervention consisted of 4 weeks of donepezil (an acetylcholinesterase inhibitor) at 5 mg/day, followed by 4 weeks at 10 mg/day, and 2 weeks of combined therapy with donepezil (10 mg/day) and intensive language action therapy. The drug effect was measured as the difference in WAB-AQ between baseline and week 8.^69^

Another trial by the same group included 20 patients who received the same donepezil regimen during the first 8 weeks, followed by 2 weeks of combined therapy (donepezil plus intensive language action therapy) with transcranial direct current stimulation or sham.^68,70^ The drug effect was again defined as the difference in WAB-AQ between baseline and week 8.

The trial by Martins and colleagues included patients with aphasia within 3 months of a left hemisphere ischemic stroke. All participants received sertraline (a selective serotonin reuptake inhibitor) at 50 mg/day and underwent 100 hours of speech and language therapy, randomized into either an intensive (10-week) or regular (50-week) schedule.^71^ Eighteen patients completed the study, and lesion data were available for six participants, equally distributed between the two groups. The study did not show a significant difference between the intensive and regular therapy arms.^71^ The drug effect was measured as the difference in WAB-AQ between baseline and week 50.

The trial by Woodhead and colleagues included 20 patients with speech comprehension impairment more than 6 months after stroke and used the CAT comprehension score as the primary outcome.^50^ This was a cross-over trial testing two concurrent interventions: phonological training using Aerobics software and pharmacological treatment with donepezil. Donepezil was administered in a double-blind, placebo-controlled design. The two cross-over groups completed four 5-week conditions: placebo alone (P1), placebo plus phonological training (P2), donepezil 5 mg (D1), and donepezil 10 mg plus phonological training (D2). The drug effect was defined as the difference between the combined D1 + D2 and P1 + P2 CAT comprehension scores.^50^

The main predictor reflected concordance between the drug tested in each trial and the neurotransmitter system with the highest ratio identified by NeuroT-Map. For donepezil studies, concordance was defined by elevated presynaptic ratios within cholinergic circuits (α4β2 nicotinic or M1 muscarinic systems). For sertraline studies, concordance was determined by the postsynaptic serotonergic ratio. ^38^

For outcome measures, improvement on the WAB-AQ was defined as an increase of ≥10 points. This threshold represents the midpoint between those used in the original trials (5 and 15 points).^69,71^ Although no universal cut-off has been established, a 10-point increase is widely considered indicative of clinically meaningful improvement.^73–75^ For CAT comprehension, any increase from baseline was classified as improvement, consistent with the criteria used in the original study.^50^ A secondary analysis applied more stringent thresholds: ≥15 points for WAB-AQ and ≥5 points for CAT comprehension.

The Fisher’s exact or the chi-square tests were computed to assess statistical significance.

### Supplementary Material

Supplementary_information.pdf

### Data and code availability

The code of the NeuroT-Map tool is available at https://github.com/Pedro-N-Alves/NeuroT-Map.

The language *f*MRI dataset is available at https://doi.org/10.17605/OSF.IO/KZWBH.

Aphasia Recovery Cohort dataset is available at https://openneuro.org/datasets/ds004884/versions/1.0.1.

Language *f*MRI cluster maps (Figure 1), maps of the predominant neurotransmitter systems contributing to white-matter tracts connecting language clusters (Figures 2a and 2c), and lesion density maps (Figures 3a, 4a and 4c) are available at https://doi.org/10.34973/pscg-yq28.

## Acknowledgments

We thank Nicola Palomero-Gallagher and colleagues at the Max Planck Institute for Psycholinguistics, the Donders Theme 1: Language & Communication community, the members of the Clinical Neuroanatomy of Language research group, the members of the Laboratório de Estudos de Linguagem, Faculdade de Medicina de Lisboa, and colleagues at the Neurology Department at Hospital de Santa Maria for their valuable discussions and clinical insights. We further acknowledge Zoe Woodhead for her assistance with access to clinical trial data, and Ahmad Beyh for his help with Figure 1.

## Authors contributions

PNA conceived the study, developed the conceptual framework, contributed to the interpretation of the findings, drafted the initial version of the manuscript, and approved the final version for publication.

IPM contributed conceptually, provided data from her clinical trial, contributed to the interpretation of the findings, critically revised the manuscript for important intellectual content, and approved the final version for publication.

CR provided clinical data, contributed to the interpretation of the findings, critically revised the manuscript for important intellectual content, and approved the final version for publication.

MLB, GD, and AL provided data from their clinical trials, contributed to the interpretation of the findings, critically revised the manuscript for important intellectual content, and approved the final version for publication.

MTdS contributed conceptually and to the interpretation of the findings, critically revised the manuscript for important intellectual content, and approved the final version for publication.

S.J.F. conceived and supervised the study, developed the conceptual framework, contributed to the interpretation of the findings, reviewed the anatomical dissections, led the preparation and writing of the manuscript, critically revised the manuscript for important intellectual content, and approved the final version for publication.

