## Supplementary Figure, Supplementary Table for "Neurotransmitter circuits of the human language network and their disruption in post-stroke aphasia"

Alves PN *et al.*

**This PDF file includes:**

Supplementary Figure 1

Supplementary Figure 2

Supplementary Figure 3

Supplementary Table 1.

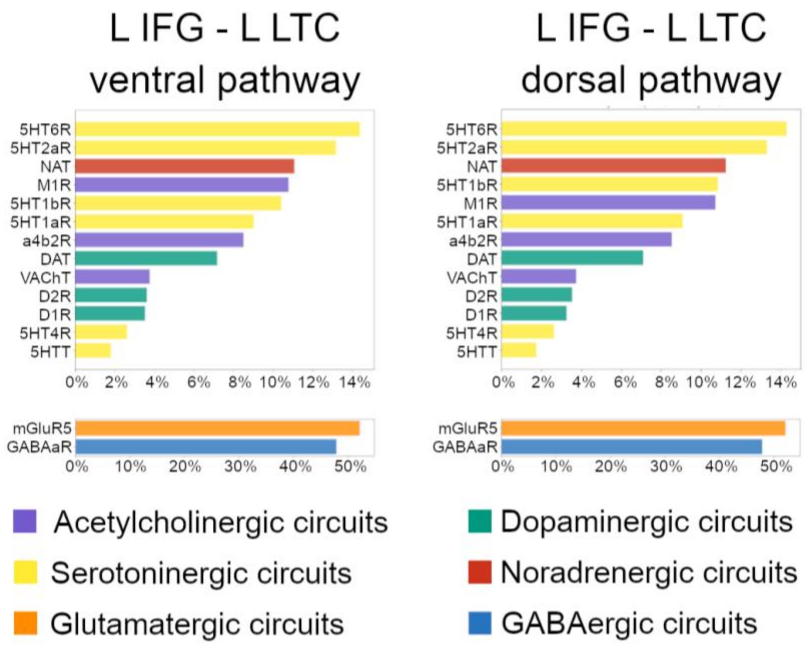


Supplementary Figure 1. Relative receptor and transporter composition of ventral and dorsal pathways between the left inferior frontal gyrus and left lateral temporal cortex. 5HT1aR, serotonin receptor 1a; 5HT1bR, serotonin receptor 1b; 5HT2aR, serotonin receptor 2a; 5HT4R, serotonin receptor 4; 5HT6R, serotonin receptor 6; 5HTT, serotonin transporter; A, anterior; a4b2R, acetylcholine receptor α4β2; D1R dopamine receptor 1, D2R dopamine receptor 2, DAT, dopamine transporter; GABAaR, GABA receptor A; L IFG, left inferior frontal gyrus; L LTC, left lateral temporal cortex; M1R, muscarinic 1 receptor; mGluR5, metabotropic glutamate receptor 5; NAT, noradrenaline transporter; VAChT, acetylcholine vesicular transporter.

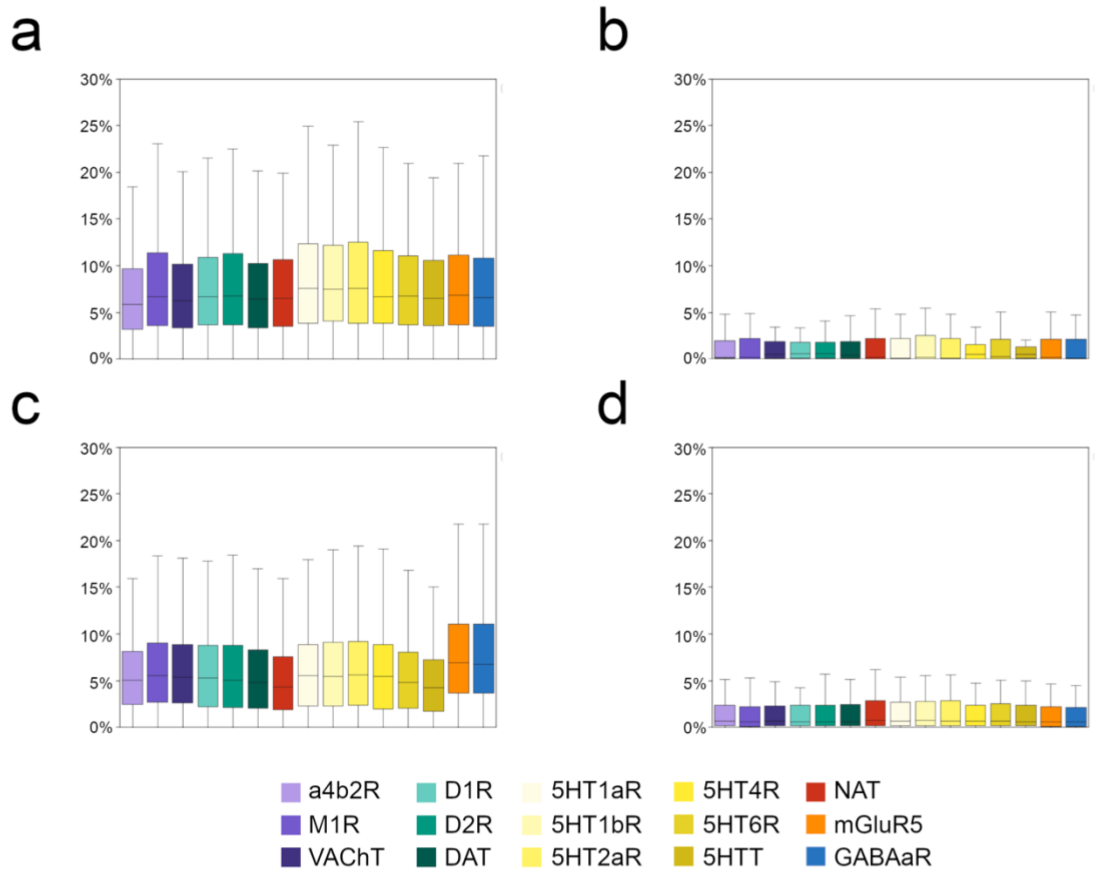


Supplementary Figure 2. Distributions of neurotransmitters' injury measures - damage to receptor and transporter density maps of patients with aphasia (a) and without aphasia (b); damage to tract maps of patients with aphasia (c) and without aphasia. 5HT1aR, serotonin receptor 1a; 5HT1bR, serotonin receptor 1b; 5HT2aR, serotonin receptor 2a; 5HT4R, serotonin receptor 4; 5HT6R, serotonin receptor 6; 5HTT, serotonin transporter; a4b2R, acetylcholine receptor α4β2; D1R dopamine receptor 1, D2R dopamine receptor 2, DAT, dopamine transporter; GABAaR, GABA receptor A; M1R, muscarinic 1 receptor; mGluR5, metabotropic glutamate receptor 5; NAT, noradrenaline transporter; VAChT, acetylcholine vesicular transporter.

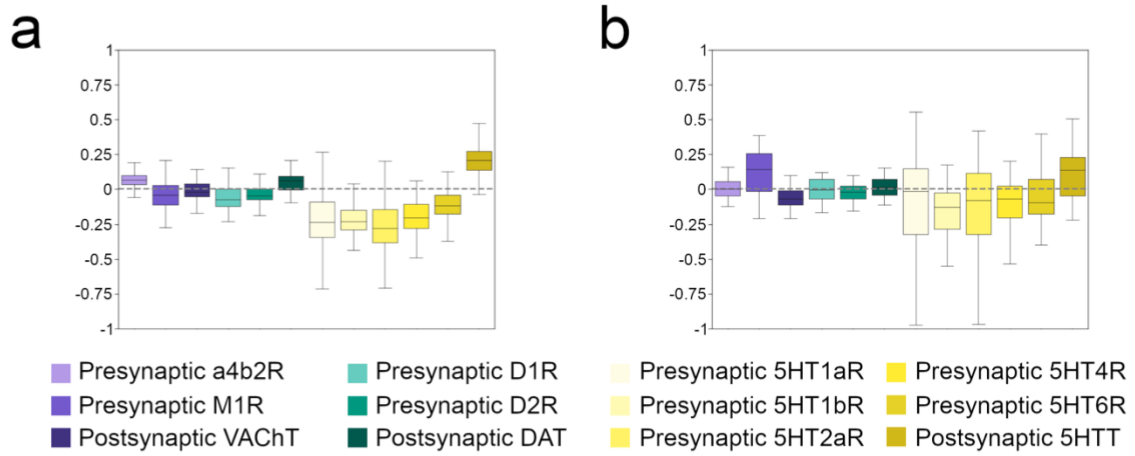


Supplementary Figure 3. Distributions of neurotransmitters' pre and postsynaptic injury ratios of patients with aphasia (a) and without aphasia (b). 5HT1aR, serotonin receptor 1a; 5HT1bR, serotonin receptor 1b; 5HT2aR, serotonin receptor 2a; 5HT4R, serotonin receptor 4; 5HT6R, serotonin receptor 6; 5HTT, serotonin transporter; a4b2R, acetylcholine receptor α4β2; D1R dopamine receptor 1, D2R dopamine receptor 2, DAT, dopamine transporter; M1R, muscarinic 1 receptor; VAChT, acetylcholine vesicular transporter.

Supplementary Table 1. Contingency table showing the relationship between drug–neurotransmitter matching (based on the maximum neurotransmitter ratio identified by NeuroT-map) and aphasia improvement, using stricter thresholds for defining aphasia improvement.

| | No aphasia improvement | Aphasia improvement |
|---|---|---|
| Drug mismatch | 38 | 3 |
| Drug match | 10 | 5 |
